# HepG2 cells with knockouts of *CYP51A1*, *DHCR24* or *SC5D* from cholesterol synthesis accumulate sterols influencing distinct regulatory pathways

**DOI:** 10.1101/2023.05.19.538399

**Authors:** Cene Skubic, Hana Trček, Petra Nassib, Andrew Walakira, Katka Pohar, Sara Petek, Tadeja Režen, Alojz Ihan, Damjana Rozman

## Abstract

Sterol intermediates of cholesterol synthesis are largely dedicated to cholesterol. Here we assess how they influence downstream gene regulatory pathways by developing knockouts (KOs) for consecutive enzymes of cholesterol synthesis in human hepatoma HepG2 cells. The KO of *CYP51*, *DHCR24*, and *SC5D* led to the build-up of specific sterols. The shared differentially expressed genes accounted for only 9% with regards to steroid metabolism and proliferation control, with majority of pathways changed in just one KO. The *CYP51* KO cells with highly elevated 24,25-dihydrolanosterol exhibited a significant increase in G2+M phase along with enhanced cancer and cell cycle pathways, likely driven by elevated LEF1 through modulation of WNT/NFKB signalling. In contrast, *SC5D* and *DHCR24* KO cells with elevated lathosterol or desmosterol, slowed cell proliferation and promoted apoptosis with downregulated E2F, mitosis, cell cycle transition, and enriched HNF1A tumor suppressor. These findings demonstrate that sterols from cholesterol synthesis control distinct gene regulatory pathways, while only early sterols can promote cell proliferation.

## INTRODUCTION

Cholesterol is by far the most abundant and studied sterol molecule, being a part of cell membranes, as well as a precursor of steroid hormones, bile acids and oxysterols, all with specific downstream signalling. Cholesterol enhances membrane fluidity and decreases membrane permeability (1–3) and is involved in the formation of lipid rafts that guide many signalling pathways (4) through interaction with proteins with cholesterol binding sites (4, 5). Other important roles are connected to cell division and proliferation, with sufficient cholesterol levels being necessary for cell cycle progression from G2 to the M phase (6, 7). Cholesterol metabolism represents an important drug targeting pathway (8, 9). Targeting this highly oxygen-dependent pathway can also alter other physiological processes (10, 11)

Much less is known about the functions of non-polar sterols from the late part of cholesterol synthesis (sterol intermediates) that are structurally similar to cholesterol (See Supp. Fig. S1 for detailed synthesis and individual sterol structures). Some of them, like desmosterol, can partially replace cholesterol in membranes (12), but not in lipid rafts (13). Increasing evidence supports specific roles of individual sterol intermediates. For example, some sterols have been proposed as ligands of the nuclear receptor Retinoic Acid Related nuclear receptor C (RORC; Zymostenol derivatives) and Liver X Receptor (LXR; desmosterol). Sterols also regulate protein degradation/activation of cholesterol homeostasis components, such as HMGCR (3-Hydroxy-3-Methylglutaryl-CoA Reductase), SREBF2 (Sterol Regulatory Element Binding Transcription Factor 2) and SQLE (Squalene Epoxidase) (14–17). They can also control T cell differentiation (18), regulate oligodendrocyte formation (19, 20), reverse protein aggregation (21) and take part in the development of hepatocellular carcinoma (22). While cholesterol is involved in the regulation of WNT signalling activities (23), the role of other sterol intermediates remains to be determined.

In humans defects in cholesterol synthesis and consequent sterol accumulation result in severe malformations, such as are Smith-Lemli-Opitz syndrome (SLOS), Antley-Bixler syndrome, Desmosterolosis, and others. Importantly, even if in all cases the same metabolic pathway (cholesterol synthesis) is disrupted, there are dramatic differences in the disease phenotypes observed, depending on which sterols accumulate in these patients (24–27).

Here we challenge the hypothesis that individual, non-polar sterol intermediates of cholesterol synthesis regulate unique sets of genes and signalling pathways that can be distinguished from their role in producing cholesterol. To investigate this, we prepared HepG2 cell models in which individual genes encoding the consecutive cholesterol synthesis enzymes CYP51A1 (Cytochrome P450 Family 51 Subfamily A Member 1), DHCR24 (24-Dehydrocholesterol Reductase), SC5D (Sterol-C5-Desaturase) were disrupted by a CRISPR-Cas9 knockout targeting strategy. After characterizing the KO cells and LC-MS/MS sterol analysis we performed a global transcriptome analysis followed by enrichment analyses of metabolic pathways and transcription factors and validated the key observations in HepG2 cells.

## RESULTS

### Targeting cholesterol synthesis

CRISPR-Cas9 generated deletion of selected genes from the late part of cholesterol synthesis (*CYP51A1, DHCR24, SCD*5), as described in Experimental procedures. HepG2 single-cell clones were initially validated at the DNA level from colonies with identical changes on both alleles (homozygous change of the gene in the DNA sequence). The detailed DNA changes, with the positions of the deletions/insertions, the guide sequences for individual gene targeted by CRISPR-Cas9 and their predicted effects on protein sequences are shown in **Supp. data, Table S1**. The selected colonies were then tested at the mRNA and protein levels (**Figure 1**). For the *CYP51* and *DHCR24* genes, an early stop codon was predicted (**Table S1**) and we were able to completely eliminate mRNA expression and consequently protein levels of CYP51A1 (**Figure 1** **A, D**) and DHCR24 (**Figure 1** **B, E**). In the case of targeting the *SC5D* gene, we introduced a 2bp deletion on both alleles, which did not result in a stop codon; consequently, mRNA and protein were still detected (**Figure 1C, F**). Nevertheless, changes in the open reading frame and modified amino acid sequence resulted in a non-functional SC5D enzyme which was proven by sterol analysis (**Figure 1** **J**).

**Figure 1.**
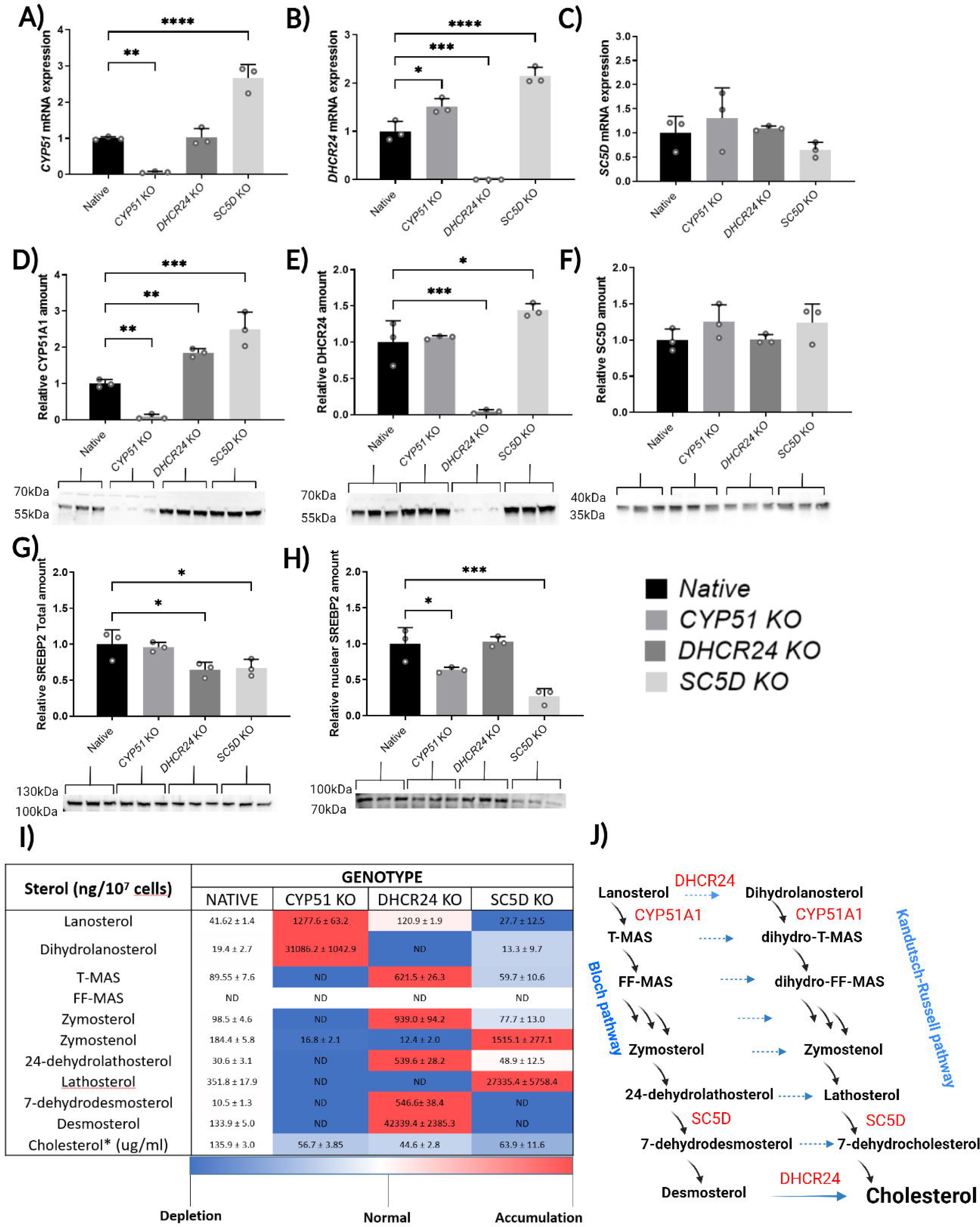
Expression of mRNA and proteins in control HepG2 cells (Native) and in knockout cells of enzymes from the late part of cholesterol synthesis (*CYP51* KO, *DHCR24* KO, *SCD* KO). Bars represent the mean + SD of three replicates from different cell culture wells. **A – C:** Relative mRNA expression of targeted genes (**A** - *CYP51*, **B -** *DHCR24*, **C** - *SC5D*). Expression data was normalized to *ACTB*, *GAPDH,* and *RPLP0* reference genes. **D** - **I**: Targeted protein expression (**D** - CYP51A1, **E -** DHCR24, **F** - SC5D, **G** c-SREBP2**, H –** n-SREBP2), by WB and appropriate antibody as described in Experimental procedures. 10 µg of proteins were loaded and normalized to total proteins. **I)** Sterol analysis by LC-MS. Sterol concentrations are represented as a mean +/- SD in ng/10^7^ cells, except for cholesterol in µg/10^7^ cells. The colours represent relative concentration compared to Native cells (Blue – Depleted; Red – Accumulated), ND – non-detectable, N=3. Cholesterol* in KO cells results from the culture medium. **J)** A simplified cholesterol synthesis, with measured sterols from the post-lanosterol part of a synthesis and the position of deleted enzymes in red. The Bloch and Kandutsch-Russell sterol pathways are indicated. For statistics, one-way ANOVA was used. *p<0.1, **p<0.05, ***p<0.01, ****p<0.001.

### Accumulation of sterols upon targeted inactivation of genes from cholesterol synthesis

Sterols successfully isolated and quantified by LC-MS/MS are represented in **Figure 1** **I)** and in **Supp. Fig. S3**, where, for each genotype, the proposed state of cholesterol synthesis is depicted. In the Native HepG2 cell line, we detected and quantified all measured sterol molecules except FF-MAS. In the *CYP51* KO cells, we detected an elevated concertation of lanosterol (1277 ng/10^7^ cells, ∼30x fold increase compared to Native) and extremely elevated concentration of 24,25-dihydrolanosterol (31 µg/10^7^ cells, ∼1500x fold change compared to Native). No other sterols were present, additionally confirming successful *CYP51* deletion. In the *DHCR24* KO, sterols from the Kandutsch-Russell pathway are absent and sterols from the Bloch synthesis pathway are accumulated (approximate fold changes compared to Native): ∼3x lanosterol, ∼7x T-MAS, ∼10x zymosterol, ∼18x 24-dehydrolathosterol, ∼50x 7-dehydrodesmosterol, ∼300x desmosterol, as the most abundant in the final step of the synthesis. The latter clearly indicates accumulation of all sterols of the Bloch pathway with higher concentrations towards the final synthesis steps. In cells with *SC5D* deletion only zymosterol (∼8x fold change) and lathosterol accumulated with ∼8x and ∼75x fold changes, respectively. Earlier sterols nor sterols from the Bloch pathway were detected in the *SCD* KO, probably due to high DHCR24 enzyme activity. Since cholesterol cannot be synthesised, the cholesterol measured **Figure 1** **I)** is from the culture medium. Cholesterol originates from cell synthesis only in Native cells where its concentration is 2-3 times higher compared to KO cell lines. Cytoplasmic and nuclear forms of SREBP2 protein are lowered, especially in the *SC5D* KO (**Figure 1** **G, H**), suggesting that disrupted cholesterol synthesis did not directly activate its synthesis pathway through SREBP2.

### Gene expression profiling, transcription factor enrichment and pathway enrichment analysis

After measuring the transcriptome and statistical analysis, differentially expressed genes (DEGs) lists (adjusted p value <0.05) for each KO cell line were obtained (full list in **Supp. materials Table S5-7**). All genes were used for the principal component analysis (PCA) plot (**Figure 2** **A**), on which the three KO cell lines cluster separately and away from the Native HepG2 cells. Compared to the HepG2 Native cell line there were 411 DEGs in *CYP51* KO, 542 in *DHCR24* KO and 765 DEGs in *SC5D* KO (**Figure 2** **B)**. Comparison of DEG lists between the three genotypes showed only 107 DEGs (or 8.9% of total) that are common to all genotypes. This was surprising since the three KO genes belong to the same synthesis pathway. Between 5-10% of genes are common to two KO lines, but over 65% of DEGs are changed just in one KO cell lines (**Figure 2** **C)**. Differences between genotypes in DEGs indicate that production of cholesterol is not the main trigger for genes being expressed, but is most likely a consequence of accumulation of specific group of sterols and their signalling. To find the differences in sterol signalling, it is important to identify genes with very high log fold change (logFC) that are differentially expressed in just one genotype. In *CYP51* KO the *CDH2* gene (N-Cadherin) is downregulated with a logFC of -3.4 (compared to logFC of -0.8 in *DHCR24 KO* and unchanged in *SC5D KO*). In the *DHCR24 KO*, *CPNE8* and *OXCT1* are highly downregulated, while *FREM1* is upregulated. In the *SC5D KO* the *ZNF385B* gene involved in the promotion of apoptosis by TP53 factor (39) is extremely upregulated with a logFC of 4.8, while unchanged in *CYP51 KO* and *DHCR24* KO. KEGG metabolic pathway analysis showed that more than half of the pathways were significantly changed just in one of the KOs (**Figure 2** **D)**. These pathways are indicated with **#** in **Figures 2** **E, F, G**. Insulin resistance, ECM-receptor interaction, NF-kappa B and PI3K-Akt signalling are modulated just in the *CYP51 KO* cell line. FoxO, Insulin and Apelin signalling are changed just in *DHCR24* KO, while Biosynthesis of unsaturated fatty acids, Fatty acid metabolism, Cellular senescence and PPAR signalling pathways are a hallmark of the *SC5D* KO cell line, with Fatty acid metabolism genes like *FADS2*, *ELOVL6*, *HSD17B12* and *CYP2J2* gene upregulated. The commonly modulated pathways changed in at least two Kos are mostly linked to disturbed cholesterol synthesis, like Steroid biosynthesis and Bile secretion.

**Figure 2.**
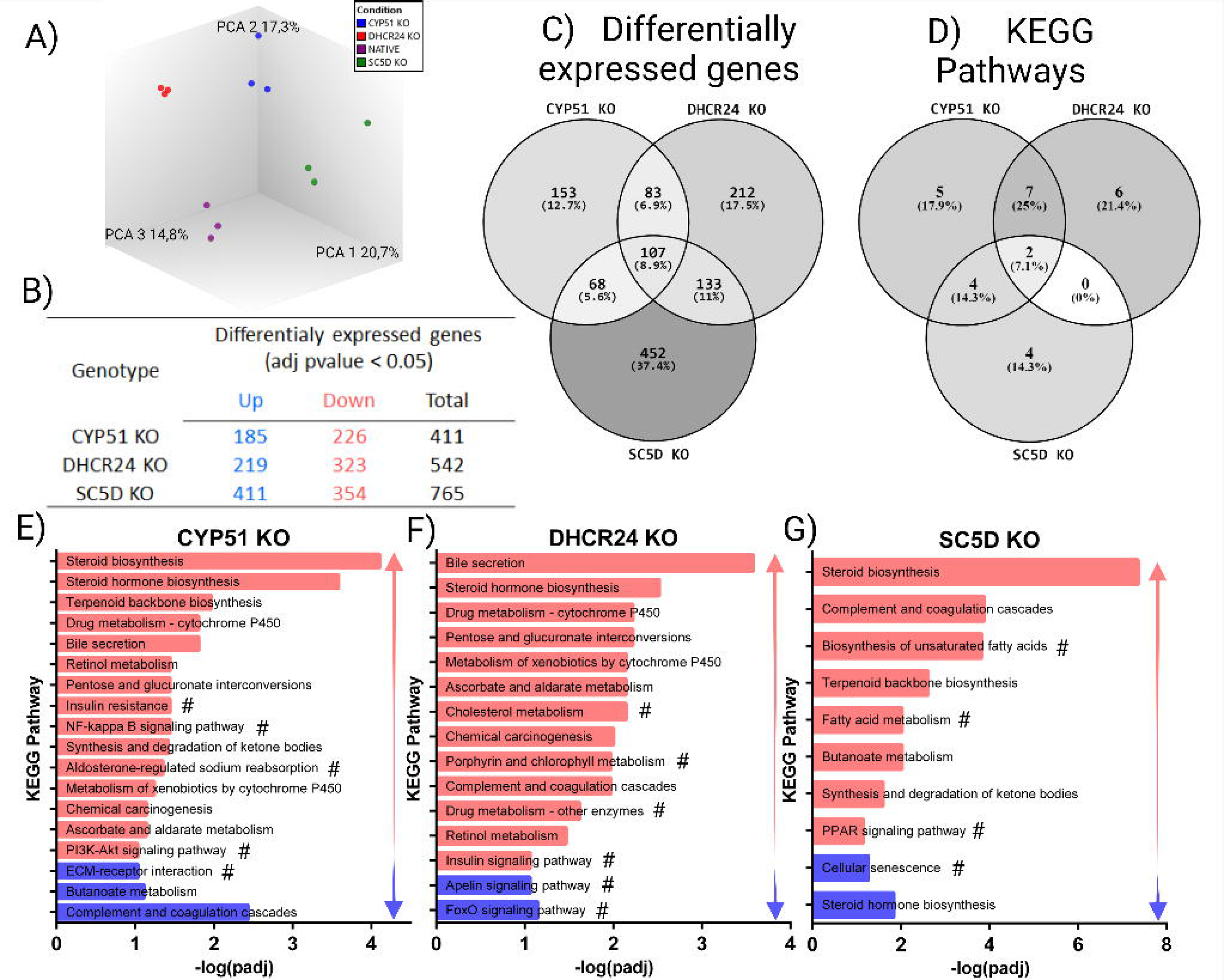
Differential gene expression (DEG) using Clarium S microarrays. **A)** PCA (Principal component analysis) of raw gene expression data for all four conditions (Native, *CYP51* KO, *DHCR24* KO, and *SC5D* KO), each measurement in three replicates from different cell culture wells. **B)** Number of up, down, and total differently expressed genes in all three genotypes compare to the Native HepG2 cell line after False discovery rate (FDR) correction (adj pvalue<0.05). **C)** Venn diagram of DEGs comparing gene expression between different genotypes. **D)** Venn diagram of changed KEGG pathways comparing different genotypes. **E)** (*CYP51* KO), **F)** (*DHCR24* KO) and **G)** (*SC5D* KO) represent volcano plots of differentially expressed genes, downregulated shown in blue and upregulated in red (FDR<0.05). The top 10 statistically significant genes are labelled. **H)** (*CYP51* KO), **I)** (*DHCR24* KO) and **J)** (*SC5D* KO) represent changes in KEGG metabolic pathways (FDR<0.05), with red showing the pathways that are upregulated and blue those that are downregulated. The # after the pathway name indicates significant change unique for this KOs cell line.

For enriched transcription factor analyses, we applied two methods that use different approaches to propose transcription factor gene regulatory networks based on differentially expressed genes and pathways: Transfac 2.0 (31) and ChEA (40). These returned different sets of TFs enriched in KOs cell lines. The top enriched TFs from both analyses are presented in **Figure 3, B and D)** (full list in **Supp. S11-S14**). Based on the ChEA analysis (**Figure 3** **A, B)** the TF with the lowest p-value in all KOs was FOXA2 (also HNF3B), which regulates bile acid metabolism and is associated with ER stress (41, 42). Nuclear receptors enriched in all KOs were: LXR (Liver X Receptor), PPARA (Peroxisome Proliferator Activated Receptor Alpha) and RXR (Retinoid X Receptor), which are known to act as heterodimers and regulate hepatic lipid homeostasis (43, 44). Sterol intermediates and oxysterols were proposed as LXR agonists, with desmosterol being the main sterol agonist, while lanosterol appears not to affect LXR activation (45–47). The *DHCR24* KO with high desmosterol concentration had the highest LXR enrichment (lowest p value). Our data also indicate activated LXR in *CYP51* KO where just lanosterol and 24,25-dihydrolanosterol are present. Based on previous studies it is unlikely that either lanosterol would activate LXR, but probably some secondary oxysterols or bile acids originating from accumulated (24,25-dihydro) lanosterol can do so.

**Figure 3.**
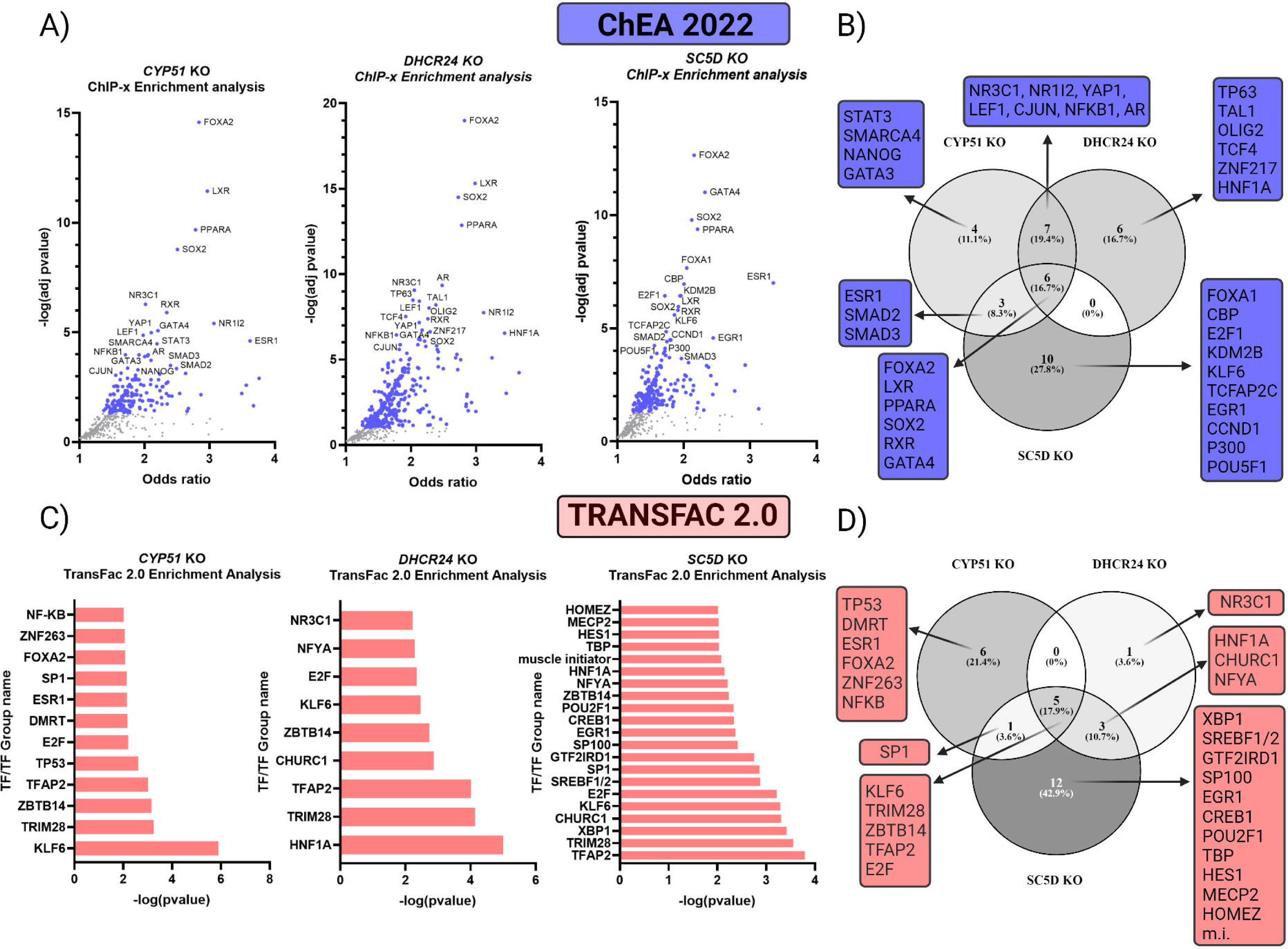
Transcription factor enrichment analysis using ChEA 2022 and TransFac 2.0. **A)** The top 20 enriched TFs from each KO cell line were used for visualization of ChEA data. **B)** The Venn diagram of enriched TFs from ChEA analysis, **C)** TFs significantly enriched in TransFac 2.0 analysis. **D)** The Venn diagram of enriched TFs from TransFac 2.0 analysis. Number of TFs are in each group is represented. % indicates the fraction of all TFs in this analysis. m.i. - muscle initiator.

Transfac analysis showed TRIM28, TFAP2A, KLF6 and E2F as the main TFs enriched in all KO cell lines with KLF6 and E2F1 also enriched in the *SC5D* KO in ChEA analysis. KLF6 (Kruppel-like factor 6) is a known tumour suppressor factor that suppress cell proliferation in a TP53-independent manner (48), while E2F1 can promote proliferation and/or apoptosis (49). Complementary to KEGG pathway analysis (**Figure 2** **E**), NFKB1 (Nuclear Factor Kappa B Subunit 1) is enriched in both datasets in the *CYP51* KO (Figure 3 A, C), which strongly indicates the potential role of 24,25-dihydrolanosterol (**Figure 1**) in the activation of the NFKB pathway. NFKB regulates the expression of genes involved in cancer development and progression, such as proliferation, migration, and apoptosis (50). The TFs enriched mainly just in the *CYP51* KO (Figure 3 B, D) are connected to proliferation control – ESR1, STAT3, NANOG and especially TP53 (51–54), together with NFKB, indicate changes in cell growth control in *CYP51* KO. LEF1 (lymphoid enhancer binding factor 1), enriched in *CYP51* KO and *DHCR24* KO, is a part of WNT signalling, connected to increased cell proliferation, epithelial-mesenchymal transition and cancer progression (55–57). Its mRNA and protein levels were highly elevated in the *CYP51* KO, as discussed later. The *CYP51* and *DHCR24* KOs also share enriched CJUN and NR1I2 (also PXR-pregnane X receptor) TFs. The TFs that were mainly enriched in the *DHCR24* KO (Figure 3 B, D) are NR3C1 (also GR-Glucocorticoid Receptor), HNF1A, TP63, and OLIG2. The latter activates the expression of myelin-associated genes that are normally expressed only in brain tissue and not in the liver. The enrichment of OLIG2 is interesting in the context of recent discoveries showing that 8,9-unsaturated sterols can drive oligodendrocyte formation and remyelination. (19). The *DHCR24* KO uniquely shows 8,9-accumulation of unsaturated sterols, such as zymosterol, TMAS, and probably other sterol molecules that were not measured in our LC-MS run (**Figure 1**), and enrichment of OLIG2, confirming the specific function of this group of sterol intermediates. With the largest number of DEGs, the *SC5D* KO has the highest number of unique enriched TFs. These are: XBP1 an ER stress-related TF active in the unfolded protein response (58), possibly resulting due to errors in SC5D protein folding; SREBF1/2 a TF responsible for regulation of genes responsible for uptake of cholesterol, fatty acids, triglycerides, and phospholipids (59); CCND1 (Cyclin D1), which controls G1/S cell phase transition (60); and EGR1 (early growth response-1) which regulates several tumour suppressor factors and inhibits cell proliferation (61). Similar results were obtained from pathway enrichment analysis **(Supp. Fig S4, S5** and **S6)**. In the *CYP51* KO (**Supp. Fig S4**) processes involved in immune response and higher cell proliferation are upregulated, whereas these pathways that enable successful cell division are downregulated in the *DHCR24* (**Supp. Fig S5)** and *SC5D* KOs **Supp. Fig S7)**.

### Sterols modulate cell proliferation and cell cycle

Based on the transcription factor gene regulatory networks and pathway analysis, we expected major changes in proliferation and cell cycle phase in KO cells. Proliferation assays were performed under three different culture medium conditions (see also **Supp. Fig. S7**) represented in **Figure 4**: classic 10% FBS **(A)**, 10% LDS + 30 µg/ml cholesterol **(B)** and 10% LDS **(C)** (LDS-Lipid Depleted Serum – sterol concentrations from both FBS and LDP are shown in **Supp. Table S4**). In FBS, which contains cholesterol and sterol intermediates, no statistically significant difference in proliferation of KO cells was observed compared to Native cells. However, *CYP51* KO cells grew faster compared to *DHCR24* (p=0.0011) and *SC5D* KOs (p=0.0026) at the 72 h time point (statistical significance not indicated on **Figure 4** **A)**. In both conditions using LDS instead of FBS (**Figure 4** **B, C)**, *DHCR24* KO and *SC5D* KO proliferation slowed significantly at 48 h and 72 h, but *CYP51 KO* cells grew at the same speed as Native cells. Especially interesting is the fact that although the KO cells cannot synthesise cholesterol, *CYP51* KO cells seem to grow normally for 72 h in culture medium with lipid depleted serum, which only has trace amount of cholesterol. This is not the case for *DHCR24* KO and *SC5D* KO, where the accumulated sterols (**Figure 1**) are structurally more similar to cholesterol (especially in the *DHCR24* KO) and can replace cholesterol to some degree (12). 24,25-dihydrolansterol and lanosterol are the first sterols in cholesterol synthesis, structurally most different from cholesterol and can in principle not substitute cholesterol in membranes. Cell cycle analysis (**Figure 4** **D)** confirms the result from the proliferation assay. Compared to Native, the *DHCR24* KO and especially the *SC5D* KO have a higher percentage of cells in G0+G1 cell phase (58% for Native cells compared to 63% for the *DHCR24* KO and 68% for the *SC5D* KO), while *CYP51* KO (57 %) does not differ from Native in the G0+G1 phase (**Figure 4** **E)**. All KOs have a statistically significantly lower percentage of cells in S phase, with the lowest in *SC5D* KO cells. Differences between KOs are again seen in G2+M, where *DHCR24* KO (22 %) and *SC5D* KO (20 %) have lower, but *CYP51* KO (28 %) a significantly higher percentage of cells in G0+M phase compared to the Native (24 %). This is consistent with the above results showing slower growth of *DHCR24* KO and *SC5D* KO cell lines. Also in concordance are the data from the transcription factor gene regulatory networks and pathway enrichment analysis (**Supp**. **Fig. S4-6),** where in *CYP51* KO proliferation pathways are upregulated while in *DHCR24* and *SC5D* KOs pathways like E2F, mitosis and regulation of cell cycle, and cell cycle transition are downregulated. In *DHCR24* and *SC5D* KOs the HNF1A, which acts as tumour suppressor (62), is enriched.

**Figure 4.**
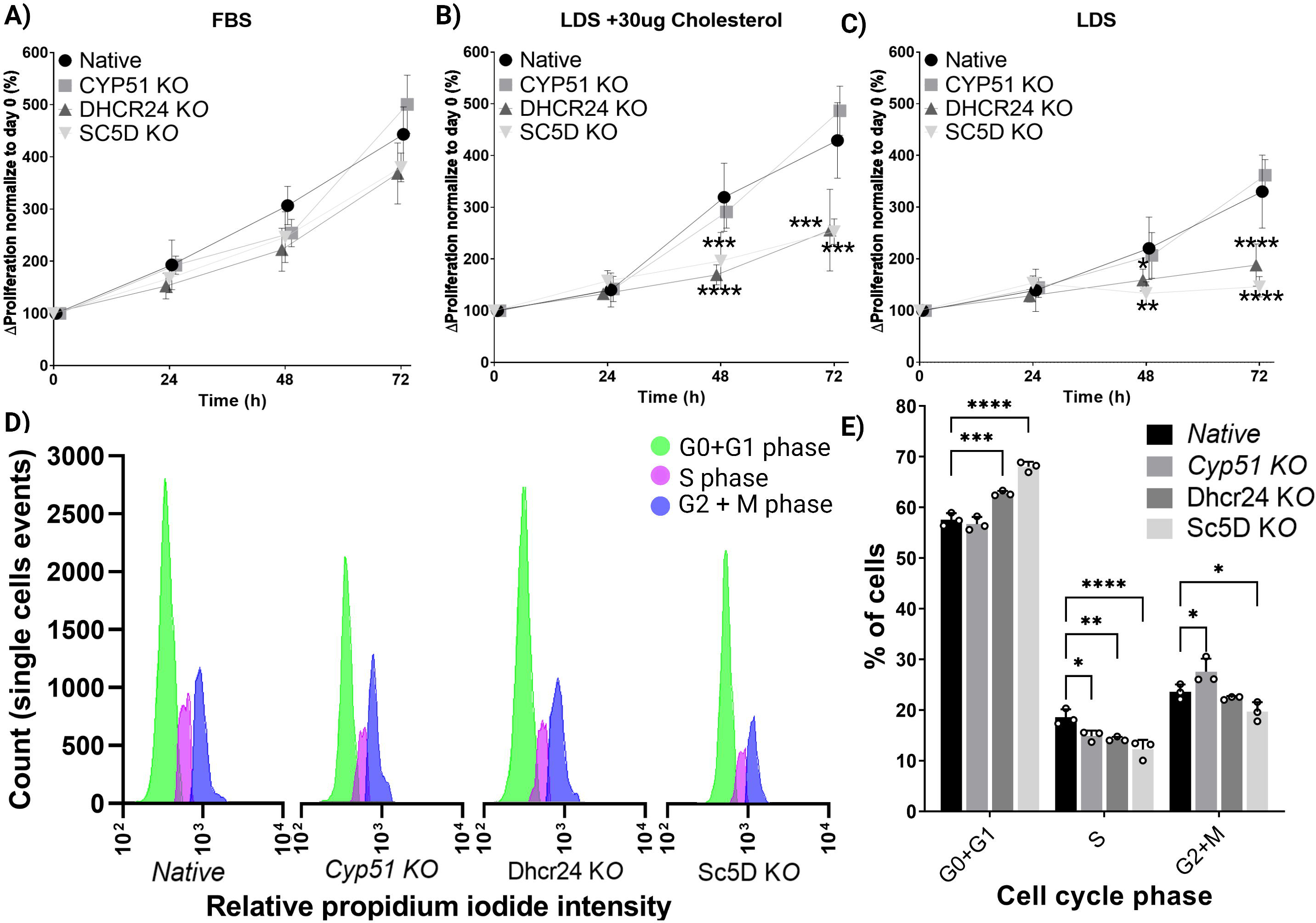
Proliferation and cell cycle analysis of HepG2 cell lines. Plots represent the proliferation of HepG2 cell lines in different serum and cholesterol conditions on **A)** FBS**, B)** LDS + 30 µg/ml cholesterol and **C) LDS**. The assay was performed using the CCK8 method and data are represented as mean +/- SE (N=6). Statistical significance was tested using One-way ANOVA with comparison to control, the Native cell line result at the same time point, p-value - *< 0.05, **< 0.01, ***< 0.001, ****<0.0001. Cell cycle measurements using propidium iodide are represented on **D)** as a direct measurement and on **E)** results normalized to total cell number (100%) and statistically evaluated. All KOs were compared with control Native cell line using the statistical test described above.

### LEF1 from WNT/NFKB pathway, but not RORC, respond to sterol-dependent signalling in HepG2 cells

Measuring the first 40 RORC target genes obtained from (63) (**Figure S10)** did not indicate changes that would depend on the sterols accumulating in specific HepG2 KO cell line. Especially desmosterol and zymosterol (14, 18) are known RORC agonists, and we expected upregulation of RORC target genes in *DHCR24* and *SC5D* KOs and downregulation in the *CYP51* KO. Using overexpression of *RORC*, immunoprecipitation and LC-MS/MS (**Supp. Fig S9**) we aimed to evaluate whether the accumulating sterols can bind to RORC protein in our cell models. However, we could not prove a specific binding.

Based on the expression profiling data and cell growth analysis we further hypothesized that sterols, especially both lanosterols, may modulate the WNT signalling pathway. **Figure 5** **A** shows differential gene expression from the WNT signalling pathway (obtained from the KEGG pathway database, PATHWAY: map04310) as detected by microarray analysis, with selected WNT genes measured by RT-qPCR (**Figure 5** **B**). Interestingly, *DKK4* is highly upregulated in all KOs and is WNT pathway antagonists (64). Another WNT antagonist is *TLE4* (65) which is highly upregulated in *SC5D* KO. The mRNA expression of *LEF1* as the key transcription factor of WNT signalling is moderately elevated in *DHCR24* KO and extremely elevated (17x fold change) in *CYP51* KO HepG2 cells, but unchanged in the *SC5D* KO. WB (**Figure 5** **C)** shows LEF1 protein detected only in *CYP51* KO, in high quantity. Transcription factors LEF1 and TCFs (T-cell factors, not changed on mRNA level, data not shown) are the main effectors of WNT signalling, whose transcription is regulated by β-catenin (CTNNB1). When the WNT pathway is active β-catenin stops being translocated to proteasomes for degradation and it is translocated to the nucleus where it binds the LEF1/TCF promoter region. N-cadherin (CDH2) whose mRNA is downregulated (logFC of -3.4) in *CYP51* KO can bind to the AXIN1-LRP6 complex and promote β-catenin degradation (66). Looking at WNT signalling proteins (**Figure 5** **C)** revealed that majority of them are unchanged. AXIN1 and cytoplasmic β-catenin were lower in *CYP51 KO*, which could indicate active state of WNT signalling, but nuclear fraction of β-catenin did not show changes. While accumulation of desmosterol, lathosterol, and other downstream sterols caused some changes in WNT genes, the LEF1 protein was not expressed. These data clearly indicate that accumulation of 24,25-dihydrolansterol caused high transcription and translation and (based on TFs enrichment) also activation of LEF1, which is known to promote cell proliferation, as seen in *CYP51* KO cells (55, 57). Recent research (67) has shown an opposite effect of sitosterol, which lowers cell proliferation through blockage of WNT-LEF1 pathway. This shows that different sterols have unique and sometimes opposite effects on LEF1 signalling.

**Figure 5.**
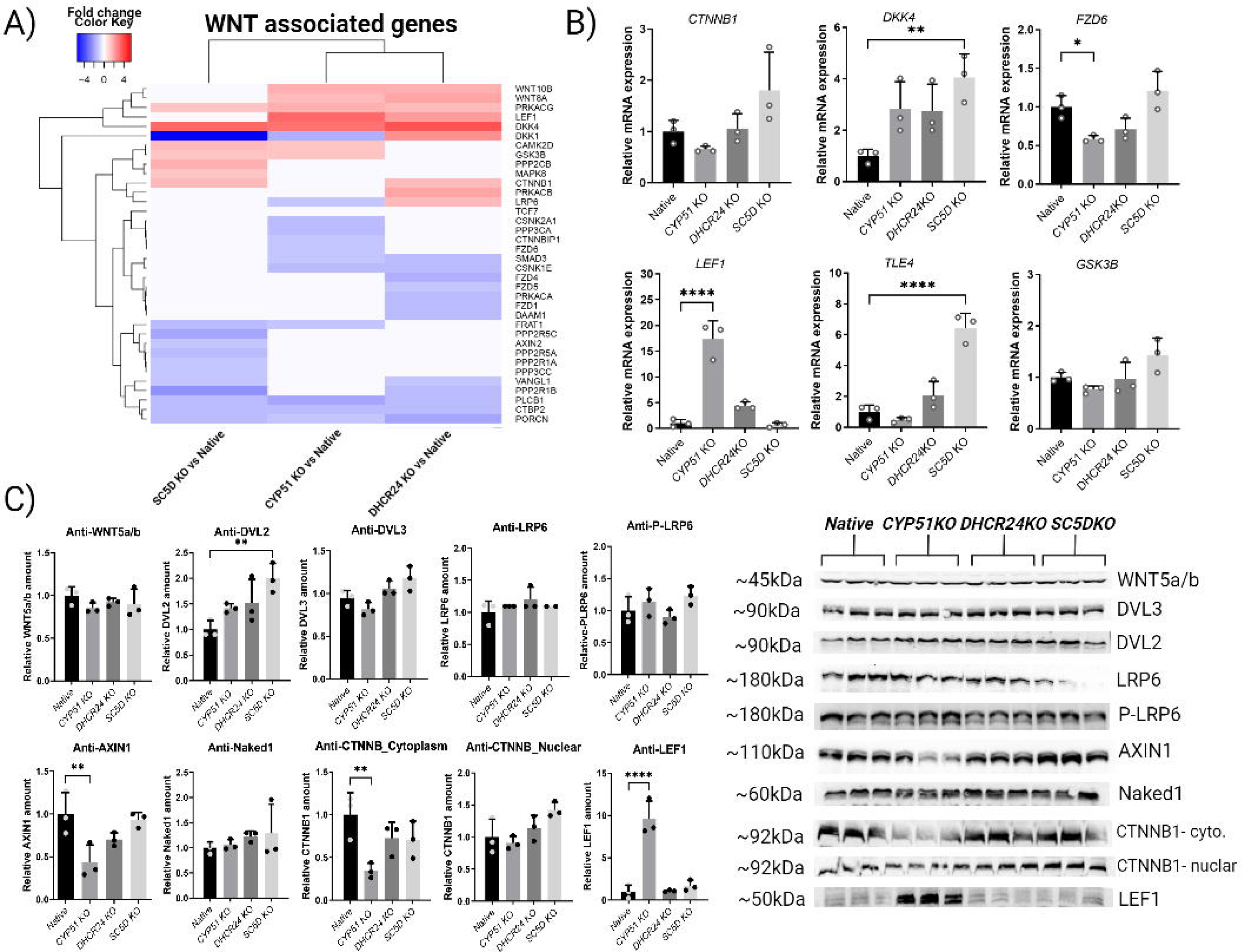
Modulation of the WNT signalling pathway. A) Heat map of genes from the WNT signalling (obtained from the KEGG database (81)), as determined by expression profiling with microarrays. White represents genes that are not changed after statistical testing, blue are downregulated and red upregulated genes, with colour intensity representing fold change. B) WNT genes measured by RT-qPCR - data are represented as mean +/- SD (three replicates from different cell culture wells). C) Expression of proteins from the WNT signalling pathway analysed by western blotting. Statistical significance was tested using One-way ANOVA with comparison to control - Native cell line. p-value - *< 0.05, **< 0.01, ***< 0.001, ****<0.0001.

Outside the WNT pathway, *LEF1* is regulated by the SMAD and NFKB TFs, both enriched in *CYP51* KO (68). The NFKB pathway regulates *LEF1* by direct binding of NFKB to the *LEF1* promoter region (68, 69). Besides *LEF1,* genes regulated by NFKB and changed in *CYP51* KO are *IL-18, ICAM-1, ICAM2 C3,* and *LYN* (Immune response group also changed in GSEA analysis **Supp. Fig**. **S4**). The third mechanism by which *LEF1* expression is controlled is by the SMAD TFs, both SMAD2 and SMAD3 are enriched in *CYP51* KO (70). There are some indications that the SMAD and WNT pathways can cooperatively regulate *LEF1* (71) and considering that WNT and NFKB can also crosstalk and regulate *LEF1* (72), changes in LEF1 resulting from 24,25-dihydrolanosterol pile-up in HepG2 cells could emerge *via* one or more of these pathways.

## DISCUSSION

Using CRISPR-Cas9, we obtained HepG2 cell lines with KOs in one of the three genes from the late part of cholesterol synthesis (73, 74) – *CYP51*, *DHCR24* and *SC5D*. This resulted in non-functional enzymes, accumulation of sterols upstream of the deleted enzymes, and depletion of downstream sterols. All KO cell lines were unable to synthesise cholesterol. In humans, the errors in the late part of cholesterol synthesis result in mental and physical defects, generally being more severe at the beginning of the post-lanosterol part of synthesis (24). In mouse models, the whole body *Cyp51* deletion was embryonically lethal (75). Deletion only in the liver resulted in lanosterol and 24,25-dihydrolansterol accumulation, which led to tumour growth and hepatocellular carcinoma-like symptoms (22, 76). Much of our findings in HepG2 *CYP51* KO cells where especially 24,25-dihydrolansterol concentration are high, align with results in the mouse models (22), where signalling pathways such as NFKB, PI3K/Akt, and the cell cycle are altered. A similar observation was recently made by Sax et al. (77), where they performed high-throughput small-molecule screens and tested oligodendrocyte formation. They observed a higher number of living cells and higher proliferation in cells treated with amorolfine, an inhibitor of CYP51A1 and EBP enzymes. This is consistent with our results, in which deletion of *CYP51* and accumulation of lanosterol and 24,25-dihydrolansterol resulted in HepG2 cells having a higher percentage of cells in G2+M and the same proliferation level as Native cells even in conditions with lower cholesterol content in the medium. Higher proliferation in *CYP51 KO* might be explained by high expression of *LEF1*, which promotes cell proliferation and has oncogene potential (56). *LEF1* can be activated through the WNT pathway, by activation of NFKB (72) or by SMADs TFs (68). Transcription was enriched in all KOs but in *SC5D* and *DHCR24* KO*s* proliferation was significantly lower and more cells were in G0 phase. In cells with higher concentration of downstream sterols, especially desmosterol in *DHCR24* KO and lathosterol in *SC5D* KO, the sterols caused slower proliferation rates and more cells accumulating in G0/1 phase. Although it is not possible to establish beyond doubt that the lower proliferation and cell cycle changes are caused by sterols downstream of zymosterol rather than by lower cholesterol levels, the depletion of which can both promote or inhibit proliferation (78, 79), the downstream sterols appear to have a direct effect on cell cycle regulation and proliferation. A possible mechanism is through TFs like factor E2F, which plays a crucial role in the cell cycle control and apoptosis, (80) and HNF1A which supresses proliferation (62). Interestingly, lathosterol accumulation in *SC5D* KO caused upregulation of fatty acid metabolism pathways and PPAR signalling. This requires clarification. Sterols from the Bloch pathway, especially desmosterol in *DHCR24* KO, caused enrichment of OLIG2, a TF usually found in the brain promoting myelination. This is interesting in the context of previously published work where it was shown that 8,9-unsaturated sterols caused remyelination of oligodendrocytes (19). From our experiments in HepG2 cells we were unable to confirm that sterols like desmosterol and zymosterol bind to RORC or activate nuclear receptor RORC as was shown previously for other cell types (14, 18). The reason could be that this sterol-RORC interaction does not occur in hepatocytes, more specifically in the HepG2 cell line, or that RORC transcriptional activity was lower compared to other transcriptomic changes caused by sterols in KO cells.

In conclusion, the well characterised HepG2 cells with knockouts of *CYP51A1*, *DHCR24* or *SC5D* are valuable novel tools to study the biochemical functions of non-polar sterols from cholesterol synthesis. Our findings, together with previous works, show that sterols from the late part of cholesterol synthesis have distinct biological functions and regulate distinct downstream gene regulatory pathways. Importantly, in HepG2 cells only early sterols can promote cell proliferation, with 24,25-dihydrolanosterol having the biggest effect through the LEF1 signalling, while sterols from the end of cholesterol synthesis suppress proliferation. Whether sterols exert their downstream effects through changing the membrane properties, or through receptor-mediated pathways, awaits to be determined. We are aware that metabolic and signalling pathways can be tissue and cell specific, so individual finding are not necessary applicable elsewhere. Further studies with focus on individual sterols in different cell types are awaited to improve the mechanistic insights.

## EXPERIMENTAL PROCEDURES

### Preparation of knock-out cells and cell culturing

HepG2 (human liver cancer cell line, Synthego, Menalo Park, CA, USA) with CRISPR-Cas9 mediated deletion of one of the enzymes from cholesterol synthesis (CYP51A1, DHCR24 and SC5D) and the Native cell line were ordered from Synthego (Synthego, Menalo Park, CA, USA) with certificate of authentication and negative mycoplasma test. Cells were provided as a mixed culture on which the CRISPR-Cas9 reaction was performed **(Suppl. Fig S2)** and were seeded 1 cell per well on a 96-well plate to obtain single cell colonies. Each colony was expanded, and the target gene region was PCR amplified and sequenced using Sanger sequencing (targeted regions are in **Suppl. Table S1**). The deletions that caused frameshifts (predicted with the Expasy translate tool (28)) were chosen, target gene expression measured using RT-qPCR (described in **3.3**), and protein expression tested by western blot using specific antibodies (See **3.4**). As a final control, sterols were isolated from knock-out cell lines and sterol intermediates quantified by LC-MS/MS as described in **3.5** and **3.6** and previously by our group (29).

### Cell culture condition

Cells were plated on 6-well plates and kept for 24 h in normal culturing conditions (37°C, 5% CO_2_) in Dulbecco’s Modified Eagle’s Media (DMEM) high glucose, with 10% FBS (Fetal Bovine Serum) and 1% P/S (Penicillin-Streptomycin, all from Sigma-Aldrich®). After 24 h, culture medium was removed, cells were washed 2 times with PBS, and DMEM with 10% lipid depleted serum (Biosera inc., France) and 1% P/S with 30 µg of cholesterol (Sigma Aldrich) dissolved in 100% ethanol was added. This cell culture medium was used to reduce the effects of sterol intermediates naturally present in FBS. After 48 h in lipid depleted medium, cells were washed with PBS and RNA isolated using Tri Reagent (Sigma Aldrich) following the commercial protocol.

### Gene expression analyses

For gene expression measurements all 4 cell lines (KOs and Native HepG2) were seeded on 6-well plates (1.5x10^5/well) and cultured as described in **3.2.** After 48 h in lipid depleted medium, cells were washed 2 times using 1x PBS. RNA was isolated using Tri Reagent (Sigma-Aldrich®) following the commercial protocol. RNA concentration and purity were measured using Nanodrop ND-1000 and RNA integrity (RIN) using Agilent 2100 Bioanalyses. Human Clariom™ S Assay (Thermo Fisher Scientific) microarrays were used to determine the transcriptome profiles. All KO lines were processed in triplicates. The expression of selected genes was measured by RT-qPCR (LightCycler 480; Roche), cDNA was prepared from 1 µg of RNA using QuantiNova Reverse Transcription Kit (Qiagen). SYBR Green I Master (Roche) was used for detection. Primer pairs for selected genes are listed in **Suppl. Table S2**.

### Protein isolation and western blot

Total proteins were isolated using RIPA buffer; nuclear and cytoplasmic proteins were isolated according to Shreiber et al. (30). All lysis buffers were supplemented with PhosSTOP and cOmplete-mini protease inhibitor cocktail (both Roche). Protein concentration was measured by a Pierce BCA Protein Assay Kit (Thermo Scientific, USA). For protein separation Mini-PROTEAN TGX Stain-Free Precast Gels (Bio-Rad Laboratories, CA, USA) were used and then transferred to Immobilon-P PVDF Membrane (Millipore, MA, USA). For total protein normalization, No-Stain Protein Labeling Reagent (Invitrogen, MA, USA) was used to stain PVDF membrane, which was then blocked using 5% milk or BSA in TBST for 1 h at room temperature. Membranes were incubated with primary antibodies (antibodies and dilutions used are represented in **Suppl. Table S3**) overnight at 4°C, washed 3 times with TBST for 5 min and incubated with secondary antibodies for 1 h at room temperature. After washing 3 times with TBST, proteins were imaged using Immobilon Classico Western HRP substrate or Immobilon Ecl Ultra Western HRP substrate (Millipore, MA, USA) on the iBright FL1500 Imaging System (Invitrogen, MA, USA) and quantified using Image J software (ImageJ, U. S. National Institutes of Health, Bethesda, Maryland, USA). All data are normalized to total protein staining.

### Sterol isolation

Sterols were isolated following the protocol described in our previous work (29). Briefly, 1x10^6^ of HepG2 cell lines (Native and 3 KO lines) were plated on a T75 culture flask for each sterol isolation (n=3 per genotype). Cells were cultured in classic DMEM medium with 10% FBS and 1% P/S. After 24 h, the cell medium was replaced with DMEM containing 10% LDS (Lipid depleted serum, Biosera inc.), 30 µg/ml cholesterol (Sigma Aldrich, Sigma Grade, ≥99%) and 1% P/S to eliminate the effect of sterol intermediates from classic FBS. After 48 h, cells were washed two times with PBS, detached using 1 mL of Trypsin (Sigma Aldrich), resuspended in 5 mL of DMEM without FBS and transferred to a 15 mL glass vial with a Polytetrafluoroethylene (PTFE) cap and cells counted for normalization. To each sample, 200 ng of internal standard Lathosterol-D7 (Avanti Polar Lipids) was added. Isolated sterols were dissolved in 300 µl of methanol and transferred to HPLC vials.

### LC-MS/MS sterol quantification

Sterol isolation and analysis was performed as described in (29). Sterols were separated on combined columns of Luna® (3 µm PFP (2) 100 Å (100 mm and 150 mm length) on a Shimadzu Nexera XR HPLC. The mobile phase consisted of 80/10/10/0.05% methanol/water/1- propanol/formic acid. HPLC was connected to a Sciex Triple Quad 3500 mass spectrometer for detection. For ionization, APCI (Atmospheric-pressure chemical ionization) was used in positive mode and detection was made in MRM (Multiple reaction monitoring) mode. The sample concentration was normalized on an internal standard (Lathosterol-d7) and sterol concentrations calculated corresponding to standards from Avanti Lipids (Avanti Polar Lipids, Alabaster, AL, USA).

### Transcriptome data analysis

The quality of the raw microarray data was assessed as follows. Firstly, microarray images were visually inspected for any visible aberrations. Further quality assessment was then done using the arrayQualityMetrics function of the arrayQualityMetrics library, and the RLE (Relative Log Expression) and NUSE (normalised unscaled standard errors) functions of the oligo library. All microarrays passed the quality checks. Array data was then processed for background correction, normalisation and summarisation using the robust multichip average algorithm (rma) in the Oligo library. Differentially expressed genes were determined by linear models using the Limma Bioconductor package (Full list in **Supp. Materials Table S8-10)**. Analysis for KEGG enrichment of gene sets was performed using the clusterProfiler library. A 5% level of significance was accepted after correcting for multiple testing using the BH-FDR (Benjamini-Hochberg False Discovery Rate) method (**Supp. Materials Table S5-7**). All analyses were performed in R v4.0.2. Data was deposited in GEO (Gene Expression Omnibus) under accession number GSE221582.

### Enrichment analyses of metabolic pathways and transcription factors

For Transcription Factor Enrichment Analysis two databases were used: TRANSFAC 2.0 (31) and ChIP-x Enrichment analysis 2022 (ChEA) (32). Release 2022.2 of the TRANSFAC MATRIX TABLE was used for TRANSFAC using input lists of differentially expressed genes for each KO cell line individually. For the background sequencing file 10.000 genes for the equivalent KO with the highest p-values were used. The TRANSFAC profile Vertebrate_non_redundant_minFP was used as a matrix, and a p-value < 0.01 was used as a cut-off for statistically significant enriched TFs. Chea analysis was done using the Enrichr online tool (33, 34) (https://maayanlab.cloud/Enrichr/). For each KO a list of differentially expressed genes (DEGs) was used as input. The top 20 TFs for each KO cell line are represented in **Figure 4** and a full list is presented in **Supp. materials Table S11** (Transfac), **S12, S13** and **S14** (ChEA).

### Pathway enrichment analysis

The GSEA 4.1.0 (Gene Set Enrichment Analysis) method was used for pathway enrichment analysis (35, 36) and results were visualized by Cytoscape software 3.8.0 (37). All parameters were used as detailed in (38). In short, a ranked gene list of all detected genes for each cell line was used as input and analysed in GSEA with 1000 permutations to obtain enriched pathways. The statistically significant (FDR<0.01) positive and negative enriched gene sets were obtained and clustered into nodes using Cytoscape and EnrichmentMap. The AutoAnotate function was used to describe the obtained clusters and then additionally manually curated. Results are illustrated in **Supp. Materials Fig. S4-6.**

### Cell cycle analysis and proliferation assays

Proliferation assays were performed using Cell Counting Kit-8 (CCK8, Dojindo Molecular Technologies, Rockville, MD, USA) according to the manufacturer’s protocol. Different culture medium conditions were tested (10% FBS, 10% LDS, 10% LDS + 30 µg/ml of cholesterol) and cells were grown for up to 72 h in 6 biological replications at 5000 cell/well on a 96-well plate. After 2 h of incubation of the reagent with cells, absorbance at 450nm was measured on an Epoch Microplate Spectrophotometer (Synergy-BioTek, USA).

Cell cycle analysis was performed on 6 well plates (1.5x10^5 cells/well) in culture conditions as described in **3.2**. After 48 h in lipid depleted medium, cells were washed with PBS, harvested by trypsinization, and fixed in 70% ice-cold ethanol at 4°C for 2 h. After fixation, cells were washed and resuspended in 300 µL propidium iodide (PI) staining solution (0.1% Triton X-100, 0.1 mg/mL RNase, 0.125 µg/mL PI) for 30 min at room temperature in the dark. Cell cycle distribution was analysed using the BD FACSCanto II system (BD Biosciences, San Jose, California, US), with the BD FACSDiva™ software (BD Biosciences, San Jose, California, US). The assay was repeated in 3 independent measurements each time in 6 technical repeats (n=6).

### Statistical analysis of WB, RT-qPCR, Cell cycle and Proliferation data

To evaluate results obtained from different cell lines, a one-way ANOVA test was used with multiple comparison against the Native cell line. The significance levels were defined as: * p < 0.05, ** p < 0.01, *** p < 0.001, and **** p < 0.0001. Unless otherwise stated results are presented as mean +/- standard deviation (error bars). Analysis and visualization were performed in GraphPad Prism 9 (GraphPad Software, San Diego, CA, USA). The statistics for microarray data, enrichment analyses and pathway enrichment analysis were performed separately as described. Venn diagrams were prepared using the Venny 2.1.0 platform (https://bioinfogp.cnb.csic.es/tools/venny/index.html).

## SUPPORTING INFORMATION

This article contains supporting information.

## DATA AVAILABILITY STATEMENT

The expression profiling data was deposited in GEO (Gene Expression Omnibus) database under accession number GSE221582.

## ABBREVIATIONS

DEGs: Differentially Expressed Genes
FDR: False Discovery Rate
GSEA: Gene Set Enrichment Analysis
KEGG: Kyoto Encyclopedia of Genes and Genomes
LDS: Lipid Depleted Serum
MRM: Multiple Reaction Monitoring
TF: Transcription Factors
WB: Western Blot

## ACKNOWLEDGMENTS

We thank John Hancock for reviewing the English language in the final manuscript.

