## Supplemental Data for "HepG2 cells with knockouts of *CYP51A1*, *DHCR24* or *SC5D* from cholesterol synthesis accumulate sterols influencing distinct regulatory pathways"

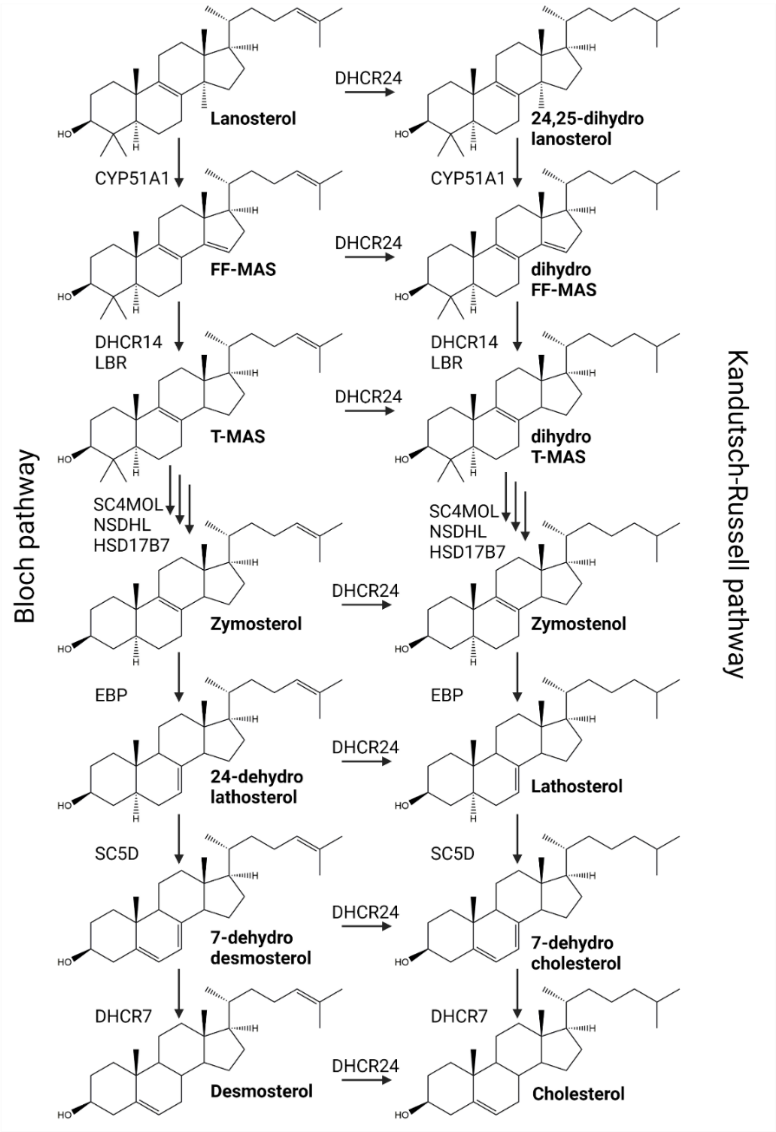


**Fig. S1.** Cholesterol synthesis from lanosterol to cholesterol. Figure represent molecular structures of sterol intermediates, their names and enzymes involved in synthesis. CYP51A1 – lanosterol 14a-demethylase; DHCR14 also TM7SF2 (Transmembrane 7 Superfamily Member 2), LBR- Lamin B Receptor , SC4MOL also MSMO-Methylsterol Monooxygenase 1 , NSDHL- NAD(P) Dependent Steroid Dehydrogenase-Like, HSD17B7- Hydroxysteroid 17-Beta Dehydrogenase 7, EBP- EBP Cholestenol Delta-Isomerase , SC5D- Sterol-C5-Desaturase , DHCR7- 7-Dehydrocholesterol Reductase , DHCR24- 24-Dehydrocholesterol Reductase.

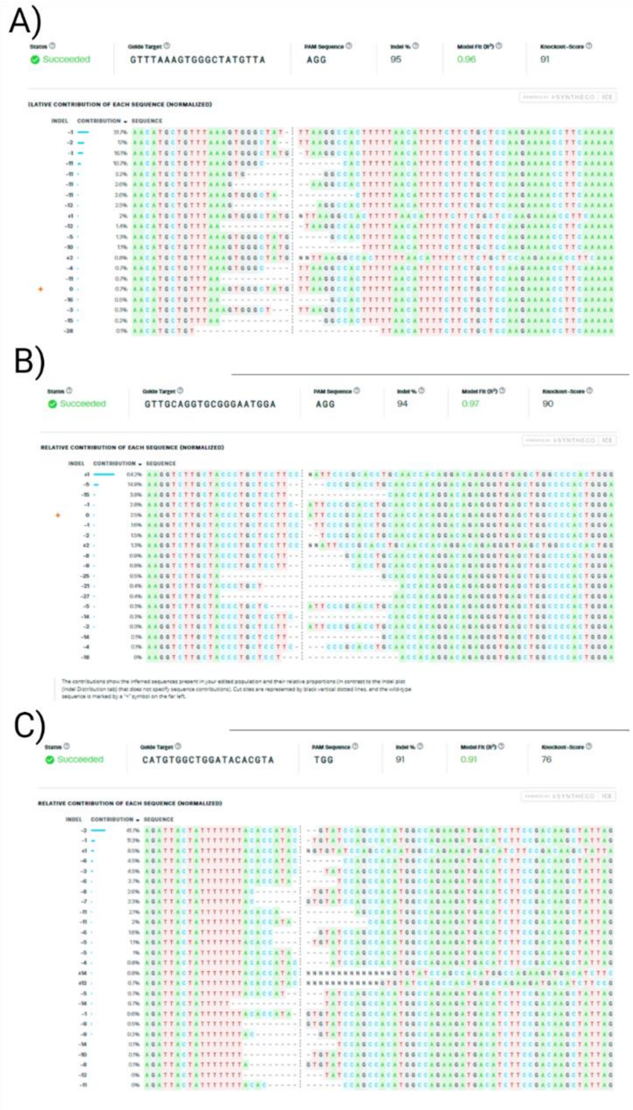


**Fig. S2.** Synthego ICE report of mixture of HepG2 cell sequenced after CRISPR-Cas9 mediated *indel* in targeted region of selected gene. A) Targeted region of *CYP51*. Guide sequence used: 5'GTTTAAAGTGGGCTATGTTA3'. B) Targeted region of *DHCR24*. Guide sequence used: 5'GTTGCAGGTGCGGGAATGGA3'. C) Targeted region of *SC5D*. Guide sequence used: 5'CATGTGGCTGGATACACGTA3'. Contribution in %, correspond to number of cells with *indel* in the mixture of targeted cells.

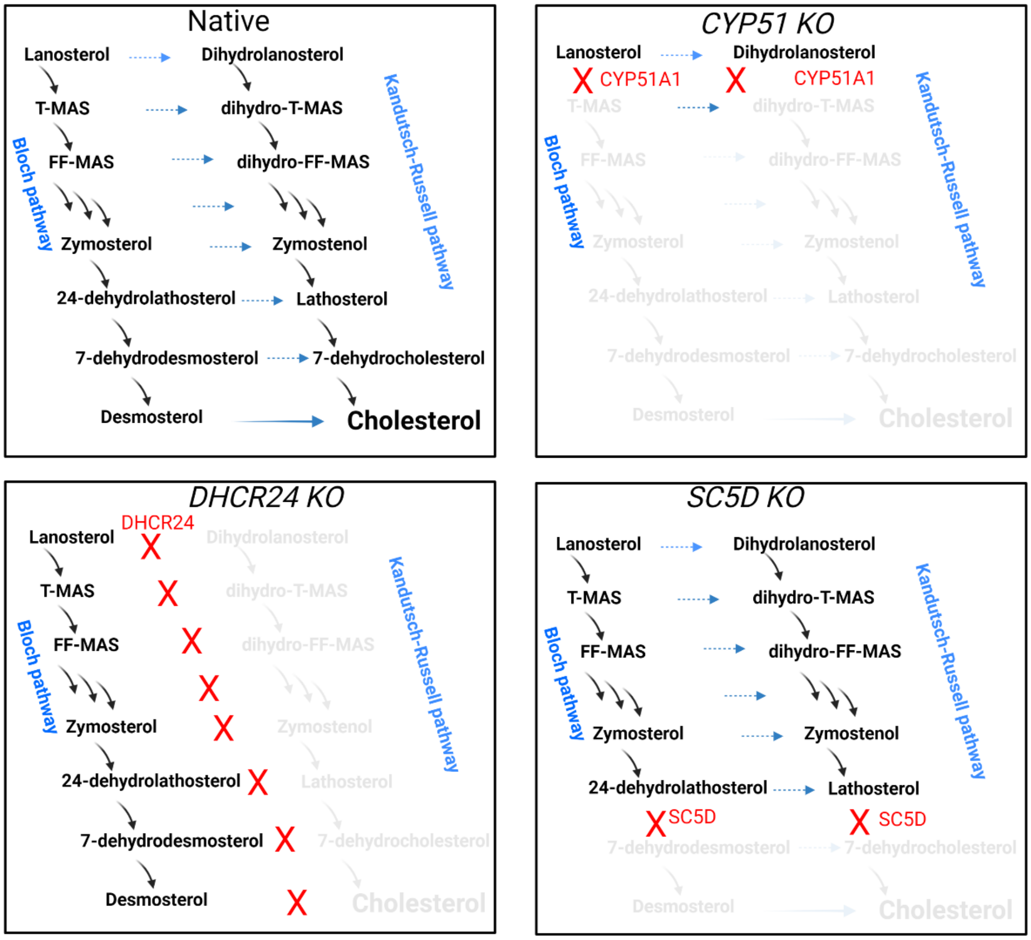


**Fig. S3.** Sterol synthesis in each HepG2 cell line used. Native represents unmodified HepG2 cells with normal cholesterol synthesis, with both Bloch and Kandutsch-Russell pathway active and end product cholesterol being synthesized. HepG2 *CYP51 KO* synthesis is stopped at the first step with two accumulating sterols. *DHCR24 KO* enriched just the Bloch pathway, most enriched are the sterols from the end of Bloch pathway. *SC5D KO* resulted in enrichment in sterols from Kandutsch-Russell pathway before the SC5D enzymatic step, with most enriched lathosterol.

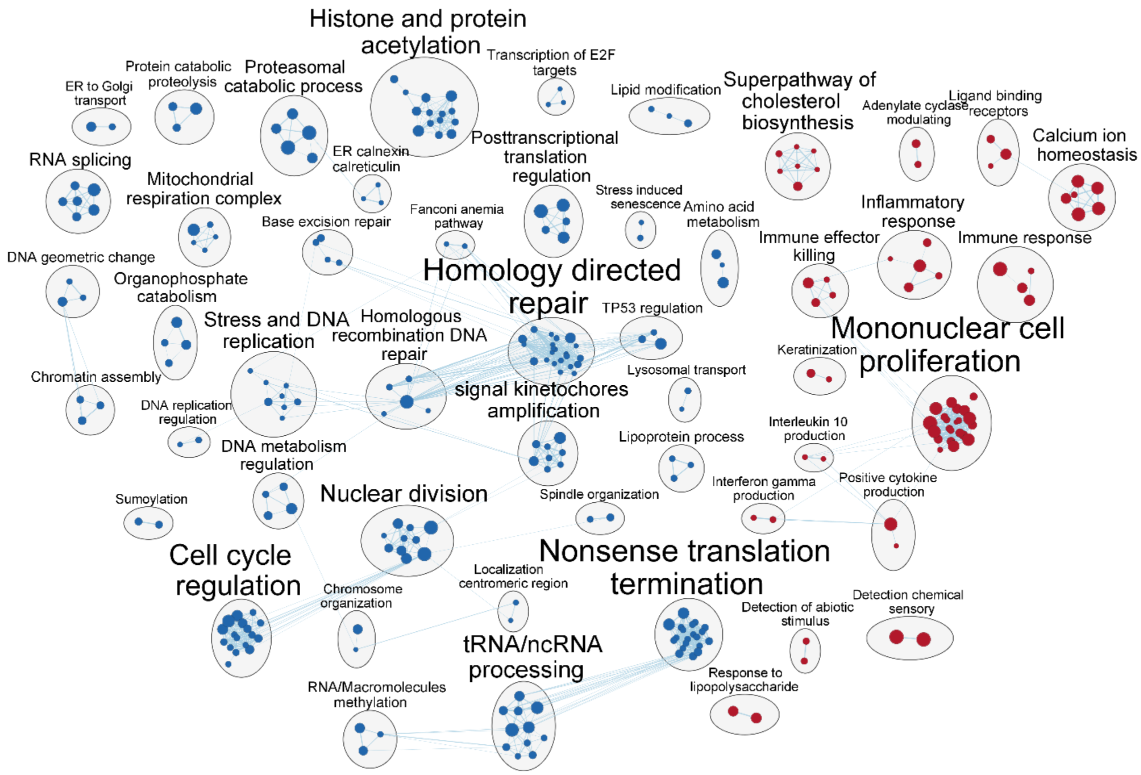


**Fig. S4.** Pathway enrichment analysis using GSEA for *CYP51 KO*. Figure created in Cytoscape based on results obtain from Pathway enrichment analysis from GSEA. Clustered pathways were auto-annotated and then additionally manually curated. In blue are downregulated pathways and in red upregulated pathways.

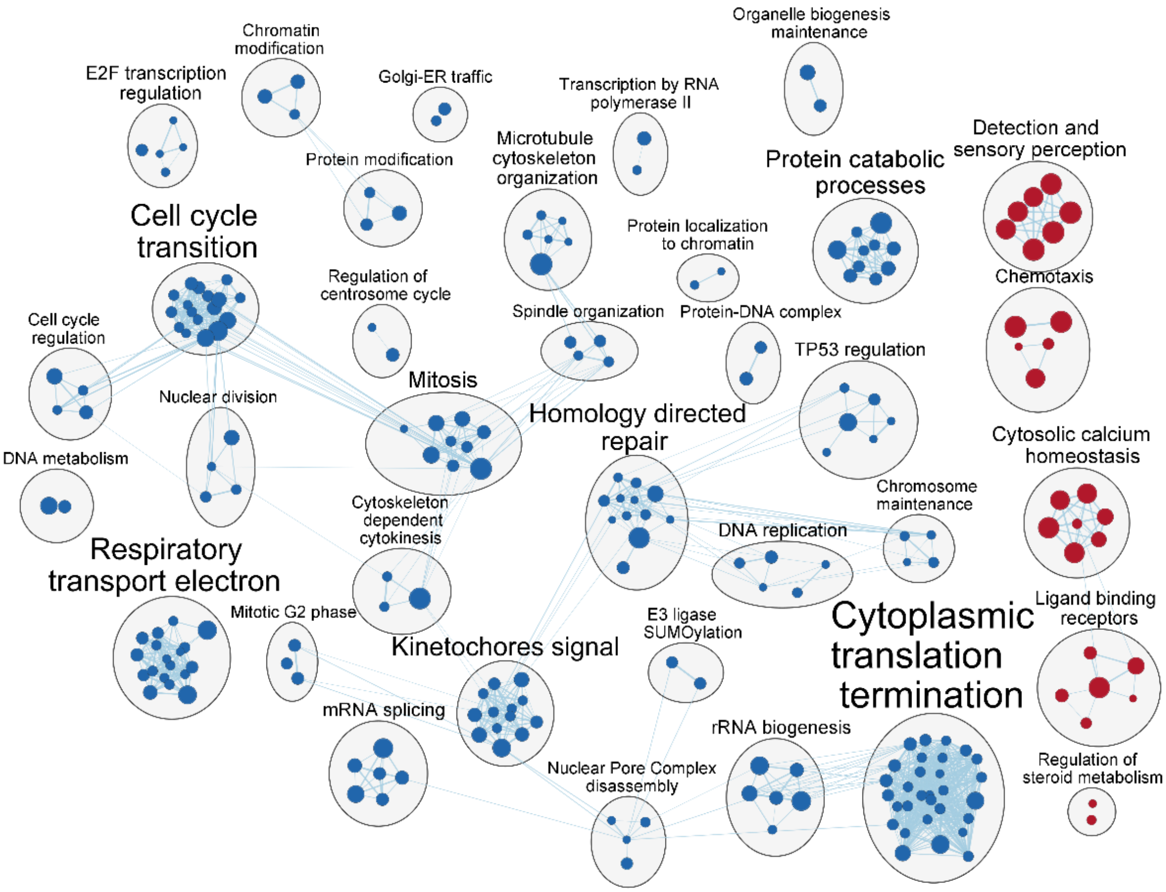


**Fig. S5.** Pathway enrichment analysis using GSEA for *DHCR24 KO*. Figure created in Cytoscape based on results obtain from Pathway enrichment analysis from GSEA. Clustered pathways were auto-annotated and then additionally manually curated. In blue are downregulated pathways and in red upregulated pathways.

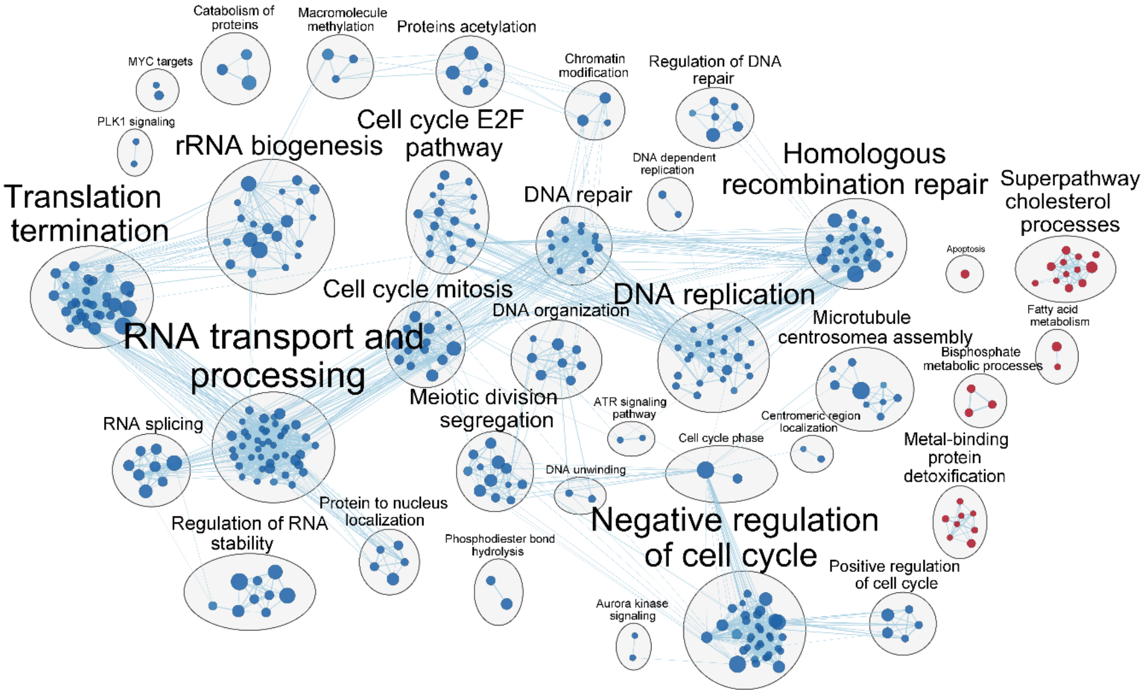


**Fig. S6.** Pathway enrichment analysis using GSEA for *SC5D KO*. Figure created in Cytoscape based on results obtain from Pathway enrichment analysis from GSEA. Clustered pathways were auto-annotated and then additionally manually curated. In blue are downregulated pathways and in red upregulated pathways.

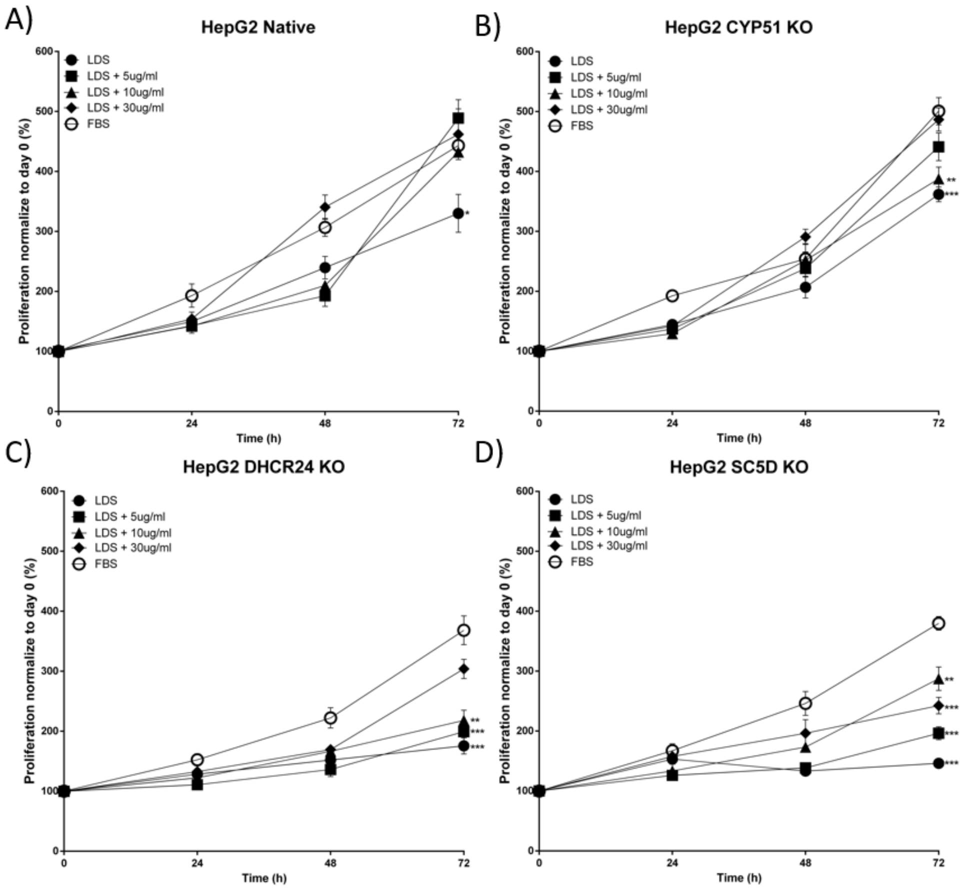


**Fig. S7.** Proliferation of HepG2 cell lines in different serum and cholesterol conditions. A) *Native*, B) *CYP51 KO*, C*) DHCR24 KO* and D*) SC5D KO*, using CCK8 assay (Cell Counting Kit-8, Dojindo Molecular Technologies, Inc.). Data are represented as mean +/- SE (n=6), statistical significance was tested by One-way ANOVA compared to control (Native HepG2 cells). P-value - *< 0.05, **< 0.01, ***< 0.001, ****<0.0001.

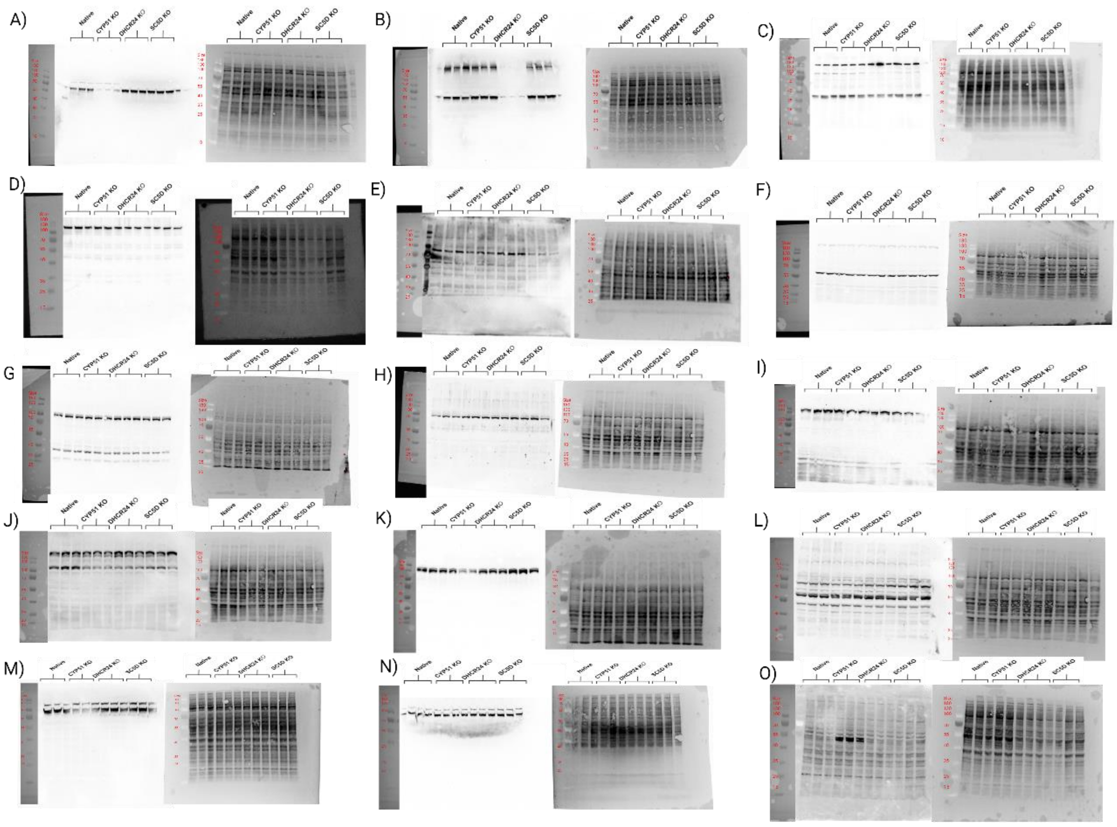


**Fig. S8.** Western blot of targeted proteins. A) CYP51A1, B) DHCR24, C) SC5D, D) SREBP2-Total proteins, E) SREBP2 – Nuclear fraction, F) WNTa/b, G) DVL3, H) DVL2, I) LRP6, J) P-LRP6, K) AXIN1, L) Naked1, M) CTNNB1 citoplasmic fraction, N) CTNNB1 nuclear fraction and O) LEF1. Whole cell lysate (10μg) from Native HepG2 cell and all of the targeted KOs was used. For total protein normalization, (right) No-Stain™ Protein Labeling Reagent (Thermo Fisher Scientific).

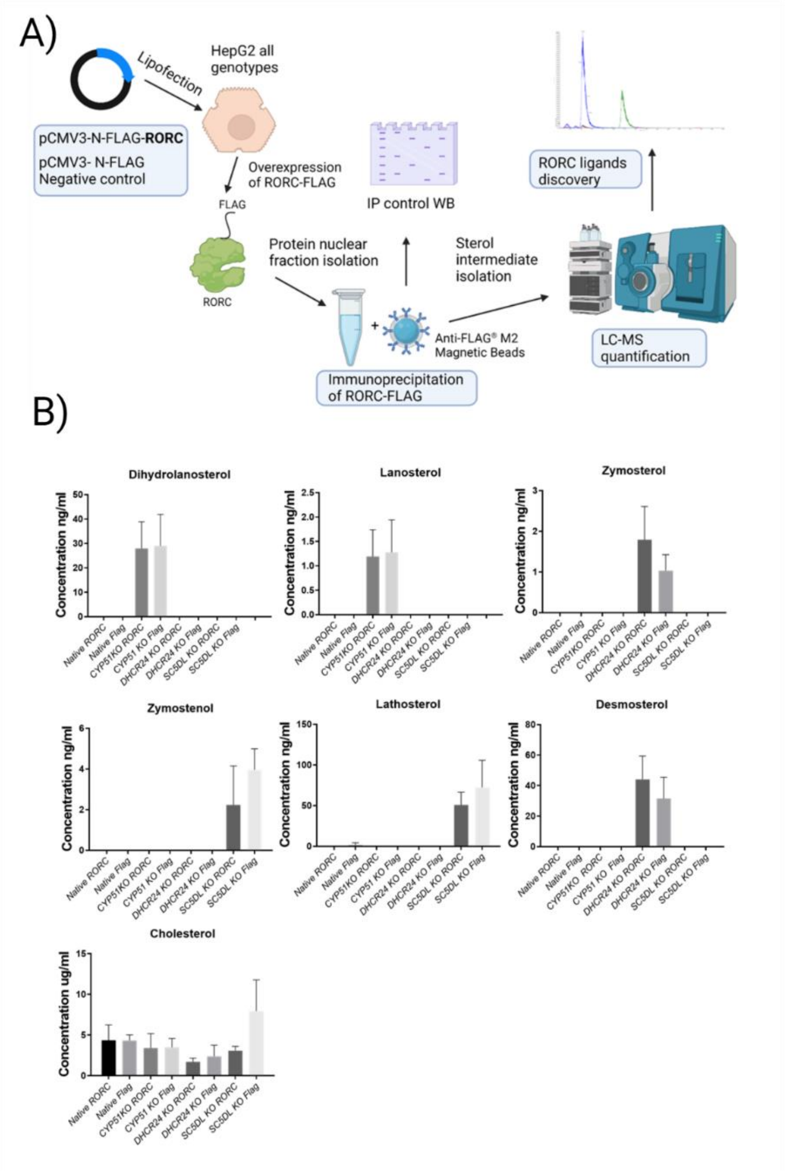


**Fig. S9.** Immuno precipitation of sterols bound to RORC. A) Schematic representation of work plan to isolate and detect sterols binding to RORC *in vivo.* B) Sterol intermediates isolated from RORC-Flag protein from one of the HepG2 cell lines. Genotype + Flag represents sterols isolated from cells overexpressing plasmid with flag tag. Genotype + RORC represents sterols isolated from cells overexpressing plasmid with RORC-Flag. HepG2 cell lines (Native and all KOs) were plated on Poly-L-lysine coated (Sigma-Aldrich) 100mm cell culture dish. Plasmids pCMV3-N-FLAG-NCV and pCMV3-N-FLAG-RORC (Sino Biological Inc.) were used and transfected to HepG2 cell lines using GenJet™ In Vitro DNA Transfection Reagent for HepG2 Cells (SignaGen Laboratories) using 7 μg of plasmid and 21 μl of reagent per plate. After 12h medium was changed with classic DMEM + 10% FBS + 1% P/S. Nuclear fraction of proteins were isolated after 48 post transfection, concentration measured using BSA. 200 μg of nuclear proteins were used for Immunoprecipitation using Anti-FLAG® M2 Magnetic Beads (Sigma-Aldrich) according to manufacturer protocol using glycine as elution. 10% of eluate was used for WB and 90% was used for sterol isolation according to protocol described in Methods 3.5. All results were normalized to input protein concentration. Similar as described in (Santori et al. Cell metab. 2015) we tried to evaluate which sterol molecules are bound to RORC in our cell models, to determine which sterols can bind and activate the RORC pathway. Unfortunately we failed to quantify specific sterols bound to RORC using Immunoprecipitation fraction of RORC protein. All quantified sterols were also present in negative controls, meaning they un-specifically co-eluted together with IP RORC, likely due to high *in vivo* concentrations. Trying to wash the IP with more organic phase (5% isopropanol) did not result in a clearer negative control (data not shown).

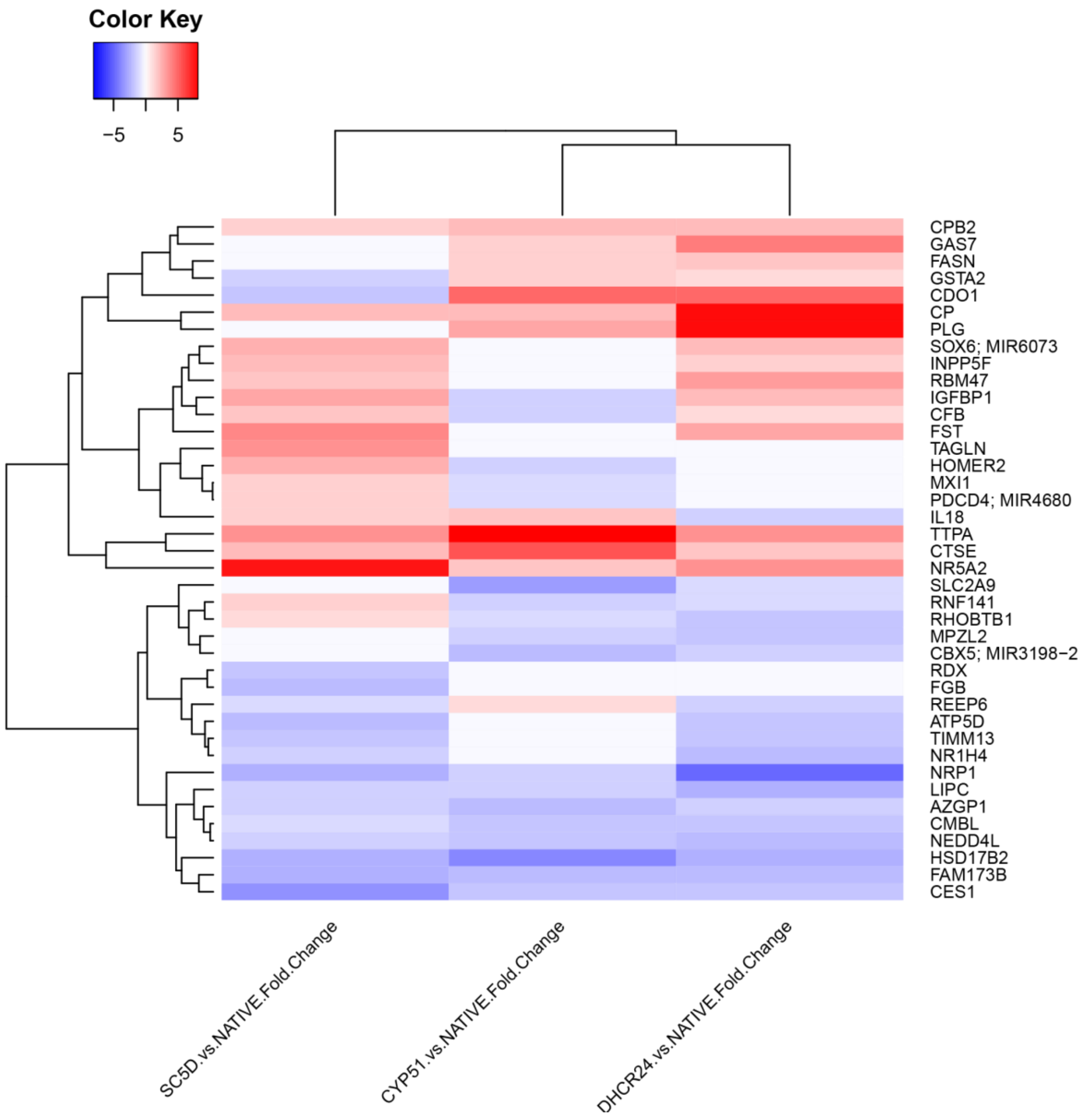


**Figure S10**: Heat map of genes, measured with microarrays, that are known targets of RORC

**Table S1.** *Indel* changes in three targeted genes in the selected single cell colonies isolated from mixture of cells targeted by CRISPR-Cas9 and their predicted effect on protein sequence. *For Sanger sequence forward primer was used.

| **Genotype** | **Targeted transcript ID** | **Guide sequence for CRISPR-CAs9 ( 5′ -3′)** | **Primer pairs for targeted region amplification and Sanger sequencing*** | **Target region of the gene** | **Allele 1 changes** | **Allele 2 changes** | **Result** | **Translate tool - ExPASy** |
|---|---|---|---|---|---|---|---|---|
| *CYP51 KO* | ENST00000003100 | GTTTAAAGTGGGCTATGTTA | FW: ACTTGCAGCCCAGTATAAAAACTG<br>RV: TGTTTCTAGTTGATGGGGAATGAGG | Exon 3 | del 2bp | del 2bp | Frame Shift | Early stop codon prediction |
| *DHCR24 KO* | ENST00000371269 | GTTGCAGGTGCGGGAATGGA | FW: GGCAGGGAGTATCTTCTATGACTG<br>RV: CACTTCGATCCTCTCTTCTAGCTC | Exon 2 | ins 1 bp | ins 1 bp | Frame Shift | Early stop codon prediction |
| *SC5D KO* | ENST00000264027 | CATGTGGCTGGATACACGTA | FW: TCTCTAACTGAGCTTTAGCATCCC<br>RV: CCATTTGTGGTTGGTCTCCAAAAT | Exon 2 | del 2bp | del 2bp | Frame Shift | No Early stop codon prediction |

**Table S2.** RT-qPCR primer list

| Gene | Forward primer (5'-3') | Reverse primer (5'-3') |
|---|---|---|
| *CYP51A1* | GCCTAATCCAGTTTTCTTGGAGC | CCACTTTCTCCCCAACTCTCA |
| *DHCR24* | GAAGCAGGTGCGGGAATGG | CATCAAGCTCAGGCAACACG |
| *SC5D* | GCCACATGGCCAGAAGATGA | TCTCTCGACGGACTTGATTCTTT |
| *ACTB* | CCAACCGCGAGAAGATGA | CCAGAGGCGTACAGGGATAG |
| *GAPDH* | AGGTCGGAGTCAACGGATTT | TGGAATTTGCCATGGGTGGA |
| *RPLP0* | TGCATCAGTACCCCATTCTATCA | AAGGTGTAATCCGTCTCCACAAGA |
| *FZD6* | GGCAGTGTATCTGAAAGTGCGC | GATGTGGAACCTTTGAGGCTGC |
| *CTNNB1* | CACAAGCAGAGTGCTGAAGGTG | GATTCCTGAGAGTCCAAAGACAG |
| *GSK3B* | CCGACTAACACCACTGGAAGCT | AGGATGGTAGCCAGAGGTGGAT |
| *TLE4* | GAACGCCATCATTGGGCAAC | GGGACTGACCTCCTAGAGCA |
| *DKK4* | ACAAGGCAGGAAGGGACAAG | GCAGACCTGTCCCTCCAAAA |
| *LEF1* | CCCGTGAAGAGCAGGCTAAA | TCTTGGACCTGTACCTGATGC |

**Table S3.** The list of antibody list used for western blots analysis

| Primary Antibody | ID | Company | Origin | Dilution | 2nd antibody used | ID | 2nd antibody Company and dilutin |
|---|---|---|---|---|---|---|---|
| **anti CYP51A1 peptide QRLKDSWAERLDFNPDRY** | anti CYP51A1 | Custom synthesised | Rabbit IgG | 1 : 1000 | Goat Anti-Rabbit IgG H&L (HRP) | ab205718 | Abcam 1 : 10000 |
| **Recombinant Anti-Seladin 1 antibody** | ab181062 | Abcam | Rabbit monoclonal | 1 : 1000 | Goat Anti-Rabbit IgG H&L (HRP) | ab205718 | Abcam 1 : 10000 |
| **Anti-SC5DL antibody** | ab221764 | Abcam | Rabbit monoclonal | 1 : 1000 | Goat Anti-Rabbit IgG H&L (HRP) | ab205718 | Abcam 1 : 10000 |
| **Anti HMGCR** | sc-271595 | Santa Cruz Biotechnology, Inc | mouse monoclonal | 1:200 | Goat anti-mouse | sc-2005 | Santa Cruz Biotechnology, 1 : 5000 |
| **Anti SREBP2** | PA1-338 | Thermo Fisher Scientific | Rabbit IgG | 1:200 | Goat Anti-Rabbit IgG H&L (HRP) | ab205718 | Abcam 1 : 5000 |
| **Wnt5a/b (C27E8) Rabbit mAb 2530** | C27E8 | Cell Signalling Technology | Rabbit IgG | 1 : 1000 | Anti-rabbit IgG, HRP-linked Antibody #7074 | 7074 | Cell Signaling Technology 1 : 2000 |

| | | | | | | | |
|---|---|---|---|---|---|---|---|
| **Dvl3 Antibody 3218** | 3218 | Cell Signalling Technology | Rabbit | 1 : 1000 | Anti-rabbit IgG, HRP-linked Antibody #7074 | 7074 | Cell Signaling Technology 1 : 2000 |
| **Dvl2 (30D2) Rabbit mAb 3224** | 3224 | Cell Signalling Technology | Rabbit | 1 : 1000 | Anti-rabbit IgG, HRP-linked Antibody #7074 | 7074 | Cell Signaling Technology 1 : 2000 |
| **LRP6 (C47E12) Rabbit mAb 3395** | 3395 | Cell Signalling Technology | Rabbit IgG | 1 : 1000 | Anti-rabbit IgG, HRP-linked Antibody #7074 | 7074 | Cell Signaling Technology 1 : 2000 |
| **Phospho-LRP6 (Ser1490) Antibody 2568** | 2568 | Cell Signalling Technology | Rabbit | 1 : 1000 | Anti-rabbit IgG, HRP-linked Antibody #7074 | 7074 | Cell Signaling Technology 1 : 2000 |

**Table S4.** Concentration of sterol intermediates measured in FBS (Fetal bovine serum) and LDS (Lipid depleted serum). Bothe samples were isolate and measured a single time, using 100µl of serum. Isolation protocol was the same as described in Methods 3.4.

| **Sterol (ng/ml)** | **FBS** | **LDS** |
|---|---|---|
| Zymosterol | 589.7 | 0.0 |
| Zymostenol | 128.2 | 0.0 |
| TMAS | 0.0 | 0.0 |
| Lathosterol | 972.7 | 0.0 |
| Lanosterol | 32.4 | 0.0 |
| 24.25-dihydrolanosterol | 25.4 | 0.0 |
| Desmosterol | 513.5 | 0.0 |
| 24-dehydrolathosterol | 35.1 | 0.0 |
| **Cholesterol (µg/ml)** | **FBS** | **LDS** |
| | 280.9 | 1.85 |

**Dataset S5 (separate file).** Table S5: List of differentially expressed genes in CYP51 KO

**Dataset S6 (separate file).** Table S6: List of differentially expressed genes in DHCR24 KO

**Dataset S7 (separate file).** Table S7: List of differentially expressed genes in SC5D KO

**Dataset S8 (separate file).** Table S8: Enriched KEGG metabolic pathways in CYP51 KO

**Dataset S9 (separate file).** Table S9: Enriched KEGG metabolic pathways in DHCR24 KO

**Dataset S10 (separate file).** Table S10: Enriched KEGG metabolic pathways in SC5D KO

**Dataset S11 (separate file).** Table S11: Transcription factors enriched using Transfac

**Dataset S12 (separate file).** Table S12: Transcription factors enriched using CHEA3 analysis in CYP51 KO

**Dataset S13 (separate file).** Table S13: Transcription factors enriched using CHEA3 analysis in DHCR24 KO

**Dataset S14 (separate file).** Table S14: Transcription factors enriched using CHEA3 analysis in SC5D KO
